# Differential CO_2_-fixation potentials and supporting roles of phagotrophy and proton pump among plankton lineages in a subtropical marginal sea

**DOI:** 10.1101/2022.02.09.479824

**Authors:** Hongfei Li, Jianwei Chen, Liying Yu, Guangyi Fan, Tangcheng Li, Ling Li, Huatao Yuan, Jingtian Wang, Cong Wang, Senjie Lin

## Abstract

Lineage-wise physiological activities of plankton communities in the ocean are important but challenging to characterize. Here we conducted whole-assemblage metatranscriptomic profiling at continental shelf and slope sites of South China Sea to investigate carbon fixation in different lineages. We catalogued 4.4 million unique genes, ∼37% being annotatable and mainly involved in microbial metabolism, photosynthesis, amino acid synthesis, oxidative phosphorylation, and two-component systems. With *RuBisCO* expression as proxy, Calvin carbon fixation (CCF) was mainly contributed by Bacillariophyta, Chlorophyta, Cyanobacteria, Haptophyta and non-diatom Stramenopiles, which was differentially affected by environmental factors among lineages. CCF exhibited positive or negative correlations with phagotrophy gene expression depending on lineages, suggesting phagotrophy enhances (Bacillariophyta, Haptophyta, and Chlorophyta) or complements (Dinophyta) CCF. Our data reveal significant potential of non-Calvin carbon fixation (NCF), mainly contributed by Flavobacteriales, Alteromonadales, Oceanospirillales and Rhodobacterales. Furthermore, in Flavobacteriales, Alteromonadales, Pelagibacterales and Rhodobacterales, NCF potential was positively correlated with proteorhodopsin expression, suggesting that NCF is energetically supported by proteorhodopsin. The novel insights into lineage-dependent potential of carbon fixation, widespread mixotrophy, and proteorhodopsin as energy source for NCF lay a methodological and informational foundation for further research to understand the carbon fixation and trophic landscape in the ocean.

**Importance:** Lineage-dependent physiologies are very important for understanding the contributions of different lineages to the biogeochemical processes in the oceanic plankton, but it is hardly possible using classical ecological methods. Even though metatranscriptomic methods have now been increasingly used to investigate physiologies of marine plankton, lineage-specific contribution to carbon fixation and phagotrophy has not received due research effort. Using whole-assemblage (prokaryotes + eukaryotes) plankton metatranscriptomic approach, with RNA quantity-based calibration to allow comparison across separately sequenced samples, this study reveals differential capacities of carbon fixation among lineages, widespread mixotrophy, and the potential of proteorhodopsin as energy source for non-photosynthetic carbon fixation. With these novel insights this study lays a methodological and informational foundation for further research to understand the carbon fixation and trophic landscape in the ocean.

## Introduction

Functional contribution of individual lineages to a marine plankton community is challenging to characterize and has been underexplored. Thanks to the rapid growth of genomic and transcriptomic databases and the increasing accessibility of metatranscriptomics, metabolic profiles in dinoflagellates[1–3], diatoms [4], raphidophytes [5] and other lineages [4] in the natural marine environment have begun to emerge. The metatranscriptomic approach has enabled interrogation of the energetic, nutritional, and defense bases underlying phytoplankton community regime shift in the process of a dinoflagellate bloom [6] and size-dependent functional differentiation in plankton communities [7]. Most of these studies have focused on one or few species in the communities, however, and a broad community perspective is limited. Carradec and colleagues reported the first global ocean atlas of eukaryotic genes based on Tara Oceans expedition [8]. Recently, a study in the tropical Pacific revealed that dinoflagellates occupy an important niche through employing numerous metabolic strategies and play a dual role in carbon transformation [9]. More functional distribution studies are needed, especially focusing on lineage-specific contributions to carbon fixation.

Calvin carbon fixation (CCF) is usually dominated by cyanobacteria in the oceanic environment and by diatoms and other eukaryotes in the coastal waters [10, 11]. Non-Calvin carbon fixation (NCF) by bacteria has been anecdotally documented or discussed [12–14], which involve five metabolic pathways in addition the Calvin-Benson-Bassham cycle and can potentially contribute as high as >30% of total oceanic carbon fixation considering that performed at night (dark C fixation) and under radiation [15]. However, NCF has not been a topic of focused study. Besides, extensive laboratory studies indicate that many planktonic eukaryotes are capable of mixotrophy, using different degrees of phagotrophy combined with complementary degrees of photoautotrophy [16, 17]. The recent study in the tropical Pacific indicates that phagotrophy of dinoflagellates is important in euphotic and the mesopelagic zones [9]. How active individual lineages are in carbon fixation, mixotrophy, and what energy source bacteria use in non-Calvin carbon fixation (NCF) in a natural assemblage are poorly understood and underexplored.

In this study, we conducted a whole-assemblage metatranscriptomic (WAM) study for a continental shelf station and a continental slope station in South China Sea (SCS). WAM profiles gene expression in eukaryotes and prokaryotes together, allowing comparison of metabolic activities across domain boundaries. We also explored RNA quantity-based calibration to reconstruct whole-assemblage metatranscriptome from separately sequenced size fractions from the same water sample. SCS is a subtropical marginal sea that connects the Tibetan Plateau and the Western Pacific Warm Pool. There are both terrestrial input from the plateau and exchange from the warm ocean currents [18, 19]. The coastal waters (within continental shelf) are influenced by several major rivers, which bring in high nutrient loads [20, 21]. The basin of the SCS (continental slope and beyond), however, is generally oligotrophic, even though it receives nutrient inputs from the seasonally fluctuated Kuroshio current, upwelling and monsoons [22, 23]. Its carbon biogeochemistry has been extensively investigated, and parts of the ocean are found to be carbon sink whereas other parts carbon source, at least seasonally [24]. However, the similarities and differences of carbon fixation characteristics between shelf and slope areas are not clear. Moreover, the microbial communities involved in biological carbon fixation and relative contribution of each major microbial group in this sea are unknown. The major objectives of our study were 1) to catalogue functional gene repertoire, 2) characterize metabolic pathways in different lineages, 3) compare capacities of Calvin carbon fixation in different lineages, 4) characterize phagotrophic activities in different lineages, and 5) assess non-Calvin carbon fixation activity and energy sources. Data revealed a high number of unigenes (4.4 million), lineage-dependent differential capacities of Calvin carbon fixation and responses to environmental conditions, major contributors of non-Calvin carbon fixation and their potential energy source.

## Results

### Retrieved genes and their functional and taxonomic distributions

The 20 samples from which WAM was analyzed were collected from 2-3 different depths in two size fractions, at two stations, hence corresponding to ten different conditions, with two biological replicates for each condition (Figure 1). Physical and chemical parameters are shown in Table S1, showing a temperature range of 18.61 to 30.25 °C, salinity of 33.62 to 34.78, N-nutrients 0.16-9.11 µM, P-nutrient 0.01-0.72 µM. WAM sequencing and bioinformatics of these samples yielded a total of 881Gb of raw reads, which were assembled into 4,499,414 unigenes with N50 of 372 bp and maximum length 71,092 bp (Table 1). Rarefaction analysis indicated that our sequencing effort was close to saturation to recover all expressed genes (Figure S1). Of these unigenes 37.23% matched genes that have functional annotations in the databases (Table S2). These unigenes were contributed by 26,269 taxa of organisms (Table S3).

**Figure 1.**
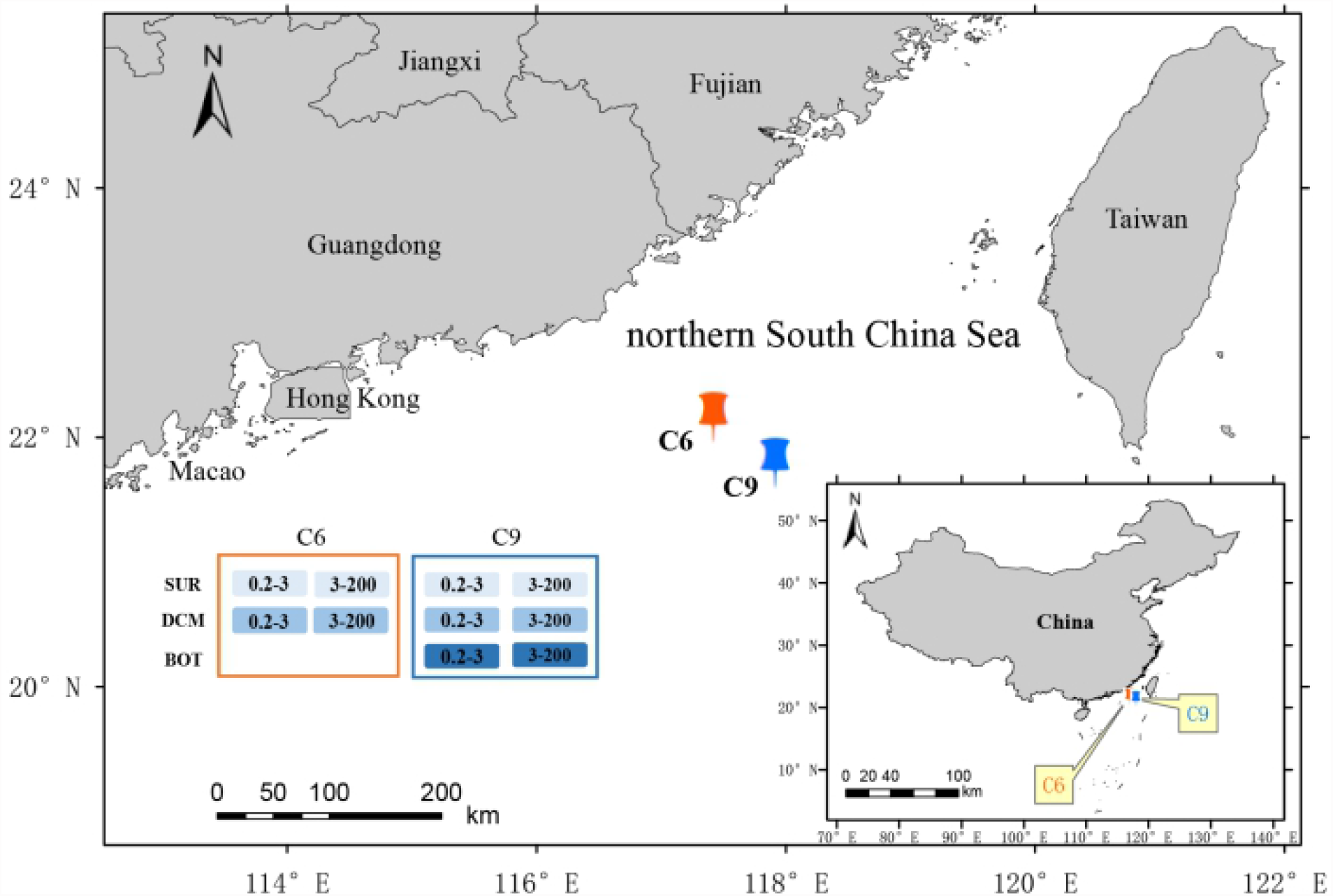
Study sites and sampling schemes. A continental shelf site (C6) and a continental slope site (C9) were sampled depths of surface (SUR) and deep chlorophyll maximum (DCM), and for C9, also the bottom of euphotic zone (BOT). For each sample, 40-50 L of water was collected and serially filtered onto 0.2–3 µm and 3–200 µm size fractions.

We focused on the top 25% most highly expression genes (HEGs) in each sample and found 35 HEGs that were common among all the samples (Table S4). These HEGs exhibited a TPM ranging 4.34-45712.73. The five genes with the highest average expression were DNA directed RNA polymerase subunit beta’, glucose-1-phosphate thymidylyltransferase, photosystem II P680 reaction center D1 protein, threonine synthase, and molecular chaperone DnaK. The 35 HEGs are mainly enriched in the five pathways of Microbial metabolism in diverse environments, Photosynthesis, Biosynthesis of amino acids, Oxidative phosphorylation, and Two-component system.

### Calvin carbon fixation (CCF) in different plankton groups and environmental effects

*RuBisCO* gene expression was used as proxy to compare the relative potential activity of CCF in plankton. Among the five-water masses, the relative activity of CCF in DCM layer was higher than SUR and BOT at both stations, with DCM of C6 station being the highest of all sample sources. The relative activity of CCF in the continental shelf area was higher than that in the continental slope area with the same water layer, and that in SUR layer of C9 station was the lowest of all sample sources (Figure 2a).

**Figure 2.**
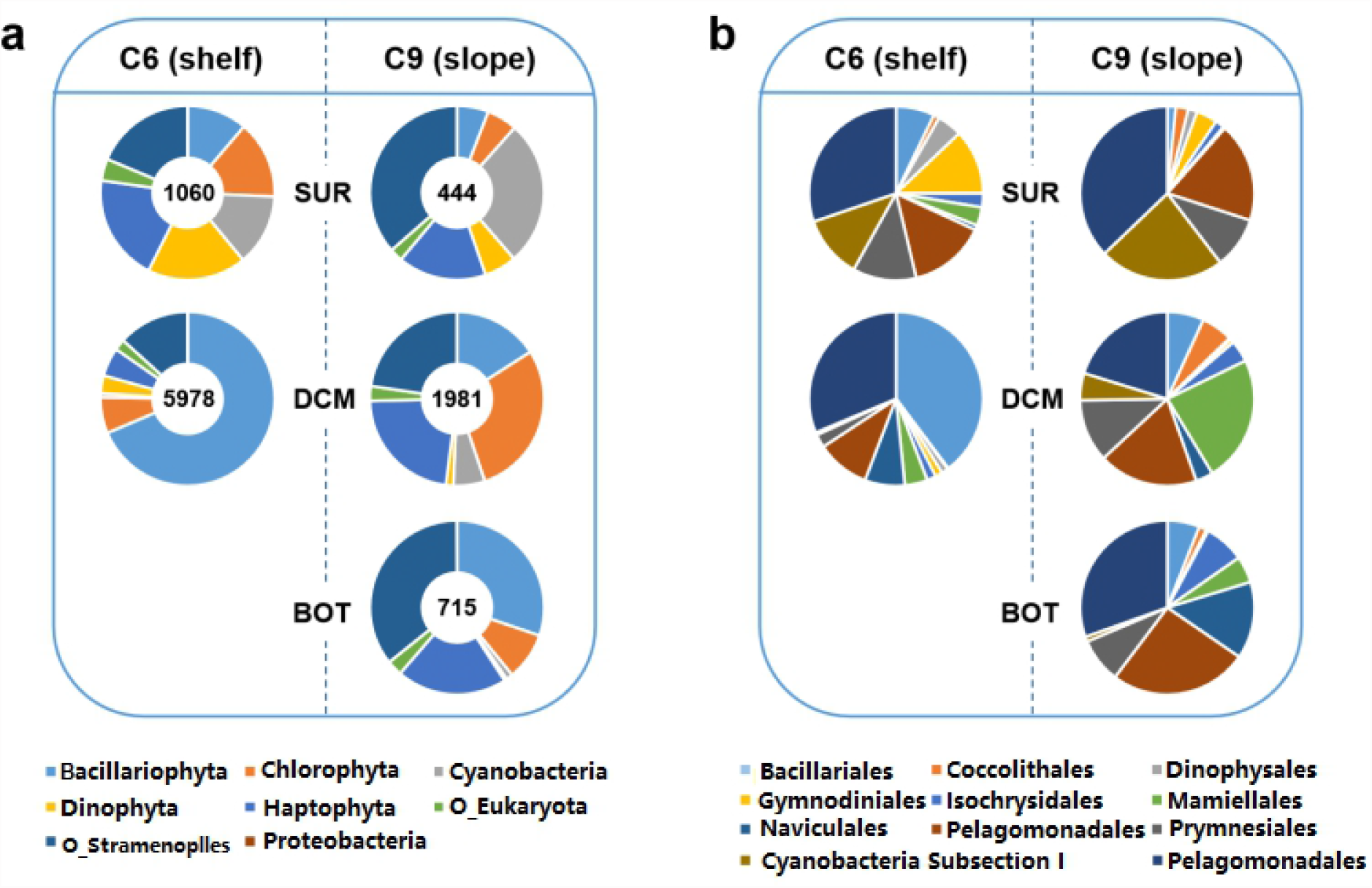
Relative contributions of major lineages to the Calvin carbon fixation (CCF) in different water layers. a, comparison at supergroup level; number in the inner circle depicts the expression level (TPM) of *RuBisCO* gene. b, comparison at order level.

At the supergroup level, the main contributors of CCF were Bacillariophyta, Haptophyta, Cyanobacteria, Chlorophyta and non-diatom Stramenopiles (O_stramenopiles) (Figure 2a). In the SUR layer at C6 station, while Proteobacteria and Other Eukaryota (O_Eukaryota) accounted for small fraction of CCF, the contributions of the other supergroups were similar, although Haptophyta contributed slightly more than did other supergroups. In the DCM layer of C6 station, the dominant CCF contributor was diatoms (Bacillariophyta), which accounted for 68.60% of total CCF potential. In the SUR layer of C9 station, the main CCF contributors were O_Stramenopiles, Cyanobacteria and Haptophyta, which accounted for 36.46%, 27.15% and 16.43% of total CCF potential, respectively. In this study, cyanobacteria contribution was the greatest in the C9_SUR environment. Bacillariophyta also contributed substantially to CCF of BOT layer at C9 station, accounting for 30.23%, and other stramenopile phyla overwhelmingly dominated the total CCF potential. For CCF-contributing Proteobacteria, only one species named Azoarcus kh32c was represented in the WAM dataset in the entire study, and its *RuBisCO* expression was detected only in the BOT layer of C9 site, contributing 0.04% to total CCF potential. At both study sites, the contribution of Bacillariophyta to the CCF of plankton increased with depth, while the contribution of Cyanobacteria and Dinophyta decreased with the increase of depth. At the same water depth, the proportion of CCF of Bacillariophyta and Dinophyta was higher at C6 station than C9 station, while the proportion of CCF of Cyanobacteria and O_Stramenopiles was higher at C9 station than C6 station. We also estimated the CCF contribution of major plankton at the order level. In total no less than 117 orders of plankton were detected to be active in CCF, among which the top ten plankton orders contributed 54.96%-76.84% of the carbon sequestration in each sample. The main orders active in CCF were Bacillariales, Pelagomonadales, Prymnesiales, Mamiellales and Naviculales (Figure 2b). The proportion of Gymnodiniales, Dinophyta, and cyanobacteria Subsection I in CCF decreased with the increase of depth, and the proportion of Naviculales in CCF increased with the increase of depth.

Further, we examined the relationship of the CCF community structure and environmental factors of different plankton at the supergroup and order levels, respectively. At the supergroup level, the CCF contribution of Bacillariophyta was negatively correlated with the concentration of P in seawater (Figure 3a). The CCF contribution of Dinophyta was positively correlated with temperature, and negatively correlated with salinity and depth. At the order level, the CCF contribution of Gymnodiniales, Isochrysidales and Naviculales was significantly related to salinity, depth, and PAR (photosynthetically active radiation), respectively, while the contribution of Pelagomonadales to CCF was positively correlated to N and Si concentrations and temperature and is negatively correlated to depth (Figure 3b).

**Figure 3.**
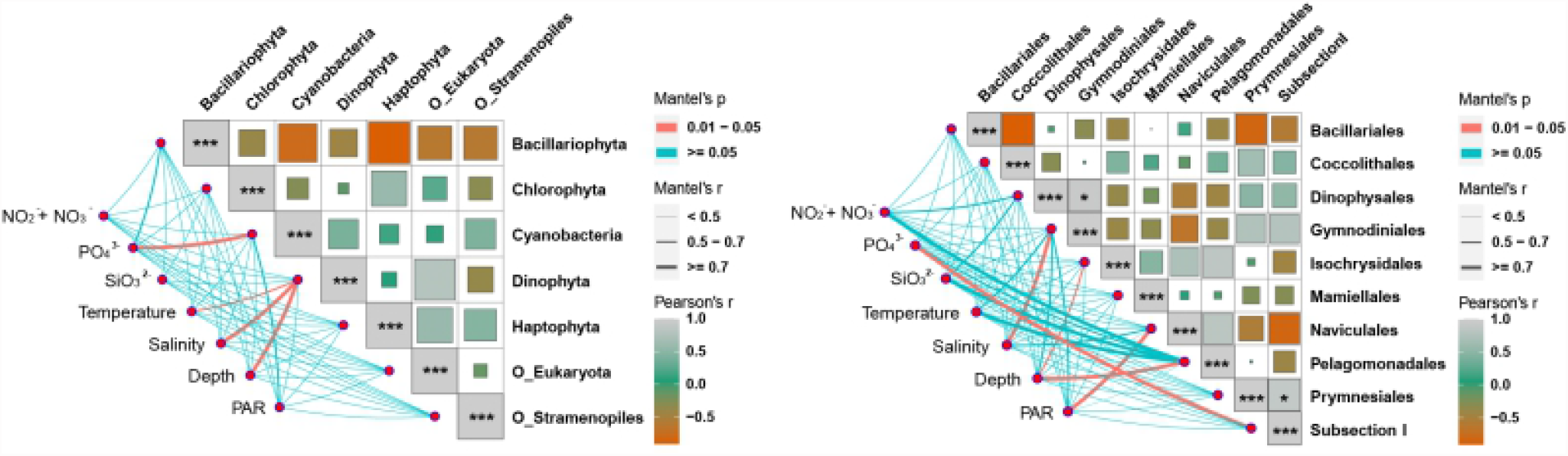
The relationship between Calvin carbon fixation (CCF) activity and environmental factors of major supergroups (a) and orders (b) of plankton. The size of the colored blocks means Pearson’s r value, and the larger the size, the larger the value. **\*\*\*P<0.01, * P<0.05**

Based on gene expression data of light-harvesting proteins and nutrient transporters (Figure S2), CCF in Dinophysales, Mamiellales and Prymnesiales was positively correlated with light energy harvesting and uptake of N and P nutrients. CCF of Bacillariales was positively correlated with acquisition of light energy, P, and Si but not with N nutrient.

### Effects of endocytosis on CCF

We analyzed the correlation between the expression of core genes in the carbon fixation pathway and the core genes of endocytosis (indicators of pinocytosis/phagotrophy) in each major CCF lineage. We found that the expression of the CCF core genes in Bacillariophyta, Chlorophyta, and Haptophyta was positively correlated with the expression of their endocytosis core genes. The correlation was negative in Dinophyta (Figure 4).

**Figure 4.**
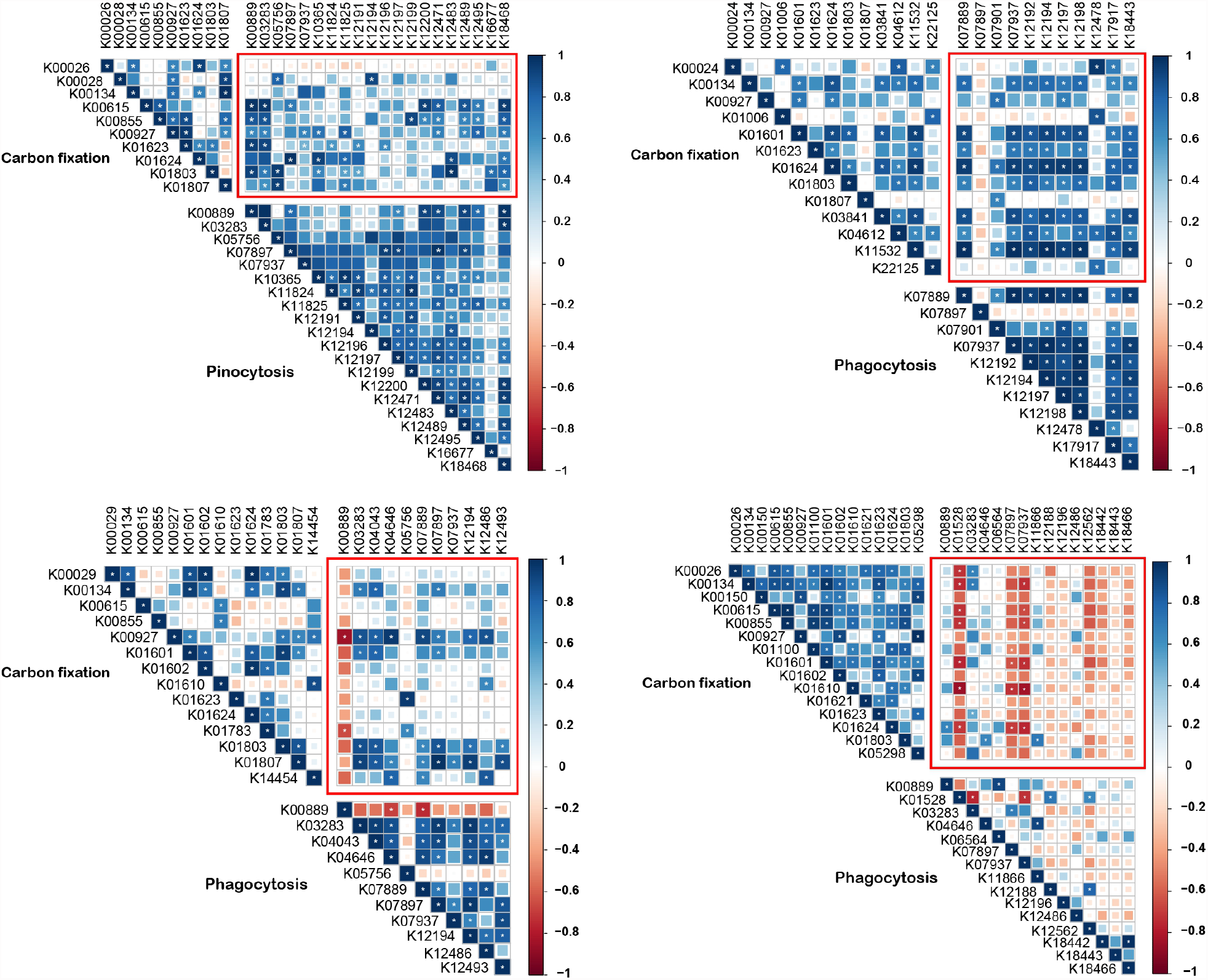
Relationship between expression of core photosynthesis genes and that of core endocytosis (pinocytosis/phagocytosis) genes in major phytoplankton phyla. The relationships are shown in red frames. Fill color of squares depicts sign and degree of correlation according to the scale bar on the right. a, Chlorophyta; b, Bacillariophyta; c, Haptophyta; d, Dinophyta. *P < 0.05. KO numbers represent genes, whose names can be found in KEGG (Table S5).

### Non-Calvin carbon fixation in different plankton groups and environmental effects

Non-Calvin carbon fixation (NCF) at both stations was contributed by Proteobacteria, Bacteroidetes, Archaea and other lineages of bacteria (O_Bacteria). The highest relative activity of NCF was in the surface layer of station C6, and the lowest was in the DCM layer at station C9 (Figure S3). At the order level, 45 orders of prokaryotes were found to express at least 15 the NCF pathway genes, among which the main five orders were Flavobacteria, Alteromonales, Oceanospirillales, Pelagibacteria and Rhodobacterales (Figure 5). Flavobacteriales contributed more than half of the NCF on the surface of the C6 station, and its contribution to NCF at the two stations decreased with increasing depth, while the NCF contribution of Alteromonadales, Oceanospirillales, Cytophagales and Nitrospinales increased with increasing depth (Figure 5).

**Figure 5.**
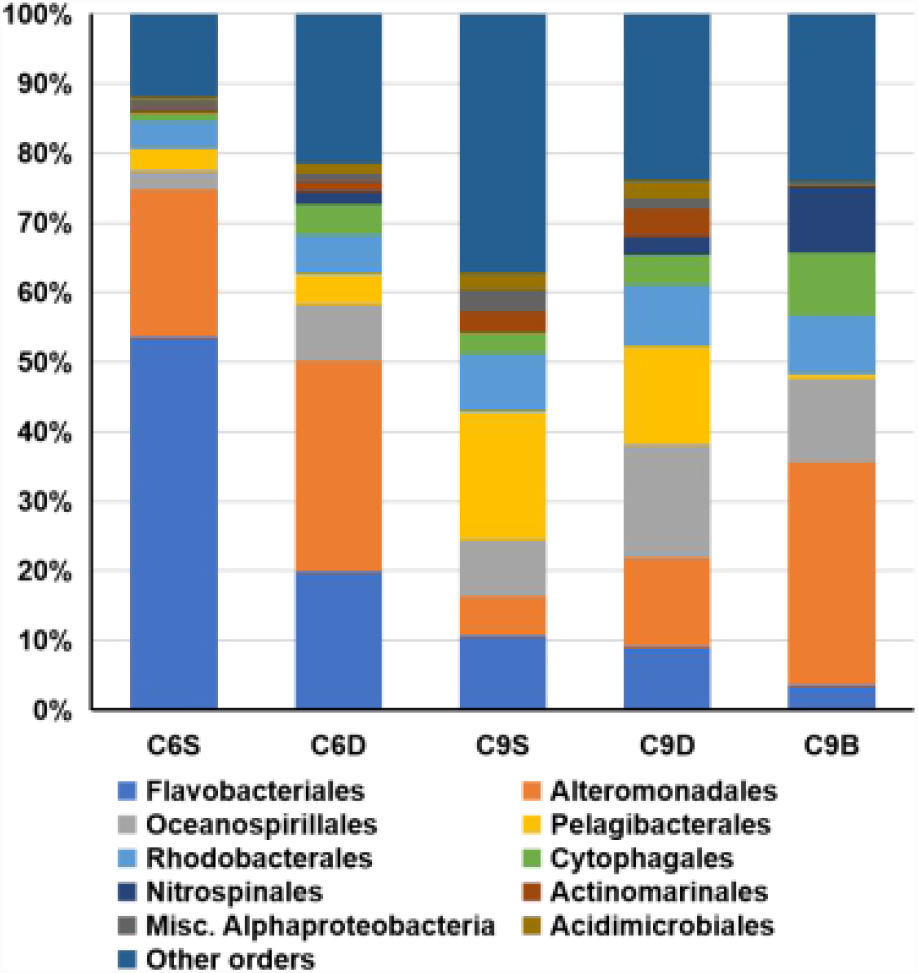
Relative contribution to non-Calvin carbon fixation (NCF) of different orders of NCF bacteria in five water masses.

We examined the correlation between NCF contribution and environmental factors of the ten major orders. We found that the NCF contribution of Misc.Alphaproteobacteria was positively correlated with temperature, depth, and N and Si concentrations. Nitrospinales’ NCF contribution was positively correlated with salinity and PAR, while Cytophagales and Oceanospirillales’ NCF contributions were affected by salinity. In addition, Pelagibacterales’ contribution to NCF appeared to be negatively affected by Si concentration (Figure 6).

**Figure 6.**
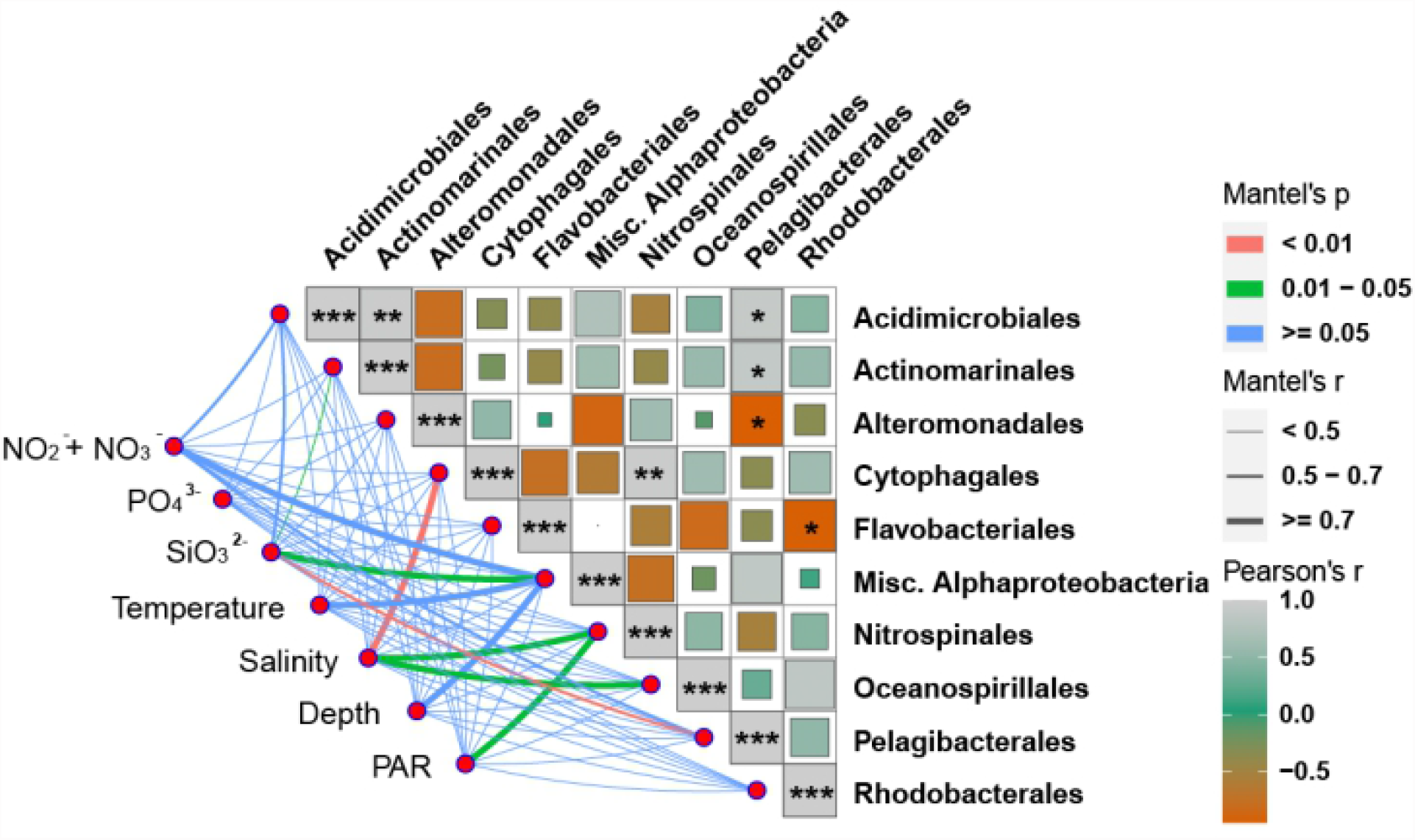
Relationship between relative contribution (%) of non-Calvin carbon fixation (NCF) by ten major orders of NCF bacteria and environmental factors. The size of the colored blocks means Pearson’s r value, and the larger the size, the larger the value. ***P<0.01, * P<0.05

In addition, by calibrating with size-fraction RNA quantities from the same water sample, we compared the NCF of two size samples in each water layer and found that both large and small sized groups of plankton contributed approximately the same (Figure S4). However, the NCF of plankton in small sized samples in SUR layer of C6 station and DCM layer of C9 station was slightly higher than that of large sized samples, and the NCF of large-sized samples in other water layers was higher than that of small-sized samples (Figure S4).

### The profile of Rhodopsin and its relationship with non-Calvin carbon fixation

A high diversity of proteorhodopsin (proton pump rhodopsin or PR) was found to be expressed at high levels in this study. In total 1,774 rhodopsin unigenes were detected, which were mainly contributed by Proteobacteria, Dinophyta, and O_Bacteria. Detailed lineage distribution and expression profiles will be reported elsewhere. Among the top six orders of prokaryotes with the highest PR expression levels, four were carbon fixers involved in NCF in this study: Flavobacteriales, Alteromonadales, Pelagibacteriales, and Rhodobacterales. Statistical analysis showed that the expression of PR was positively correlated with the expression of carbon fixation genes, with R^2^ values ranging from 0.68 to 0.90 (Figure 7). Furthermore, we used the same method to analyze 187 samples collected from 66 stations in Tara Ocean database [25] and found that the activity of non-Calvin carbon fixation of Flavobacteria in these samples was significantly correlated with its PR expression level (Figure 8).

**Figure 7.**
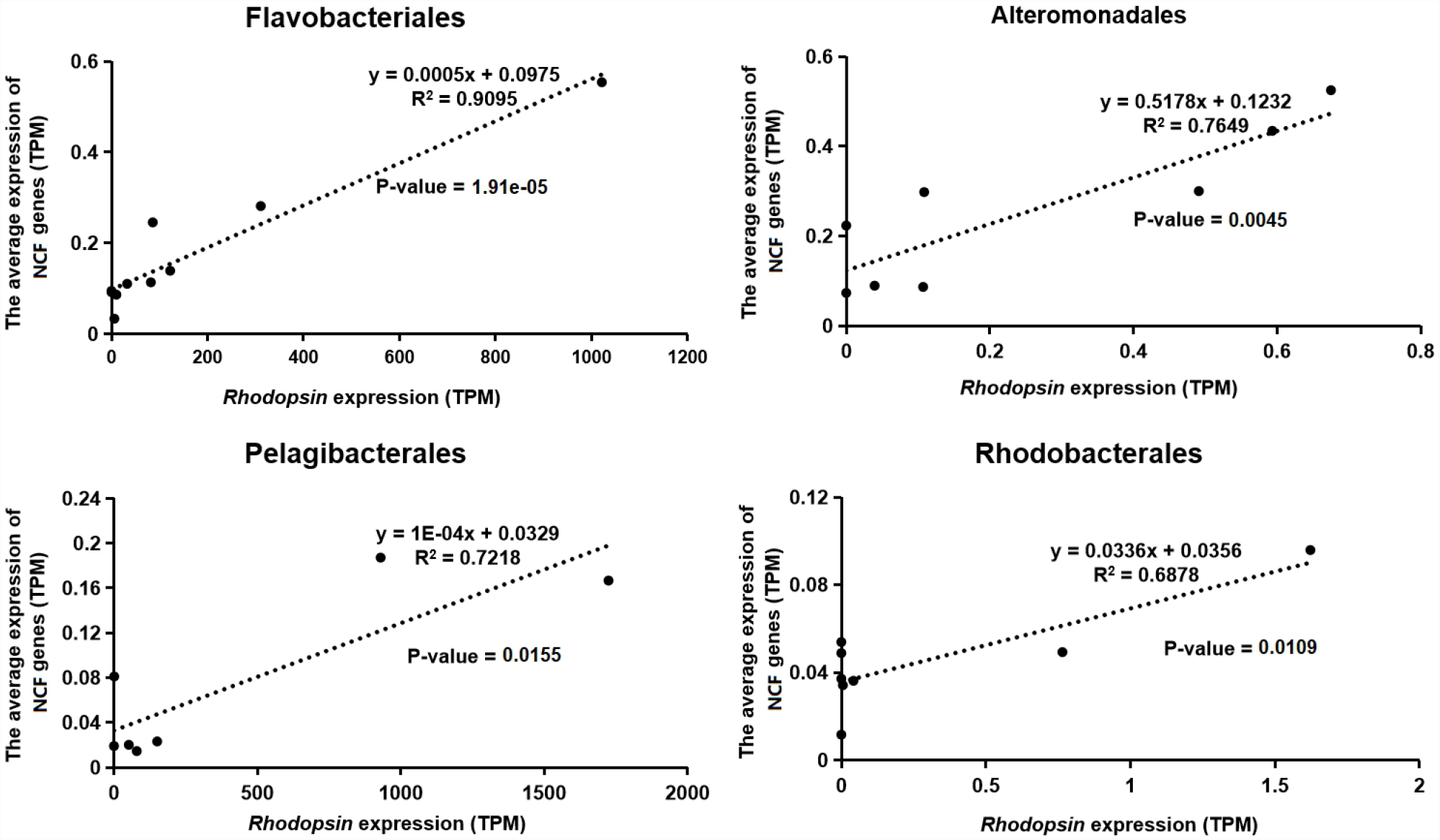
The correlation between the relative activity of non-Calvin carbon fixation (NCF) and the expression of rhodopsin in four major orders of NCF bacteria.

**Figure 8.**
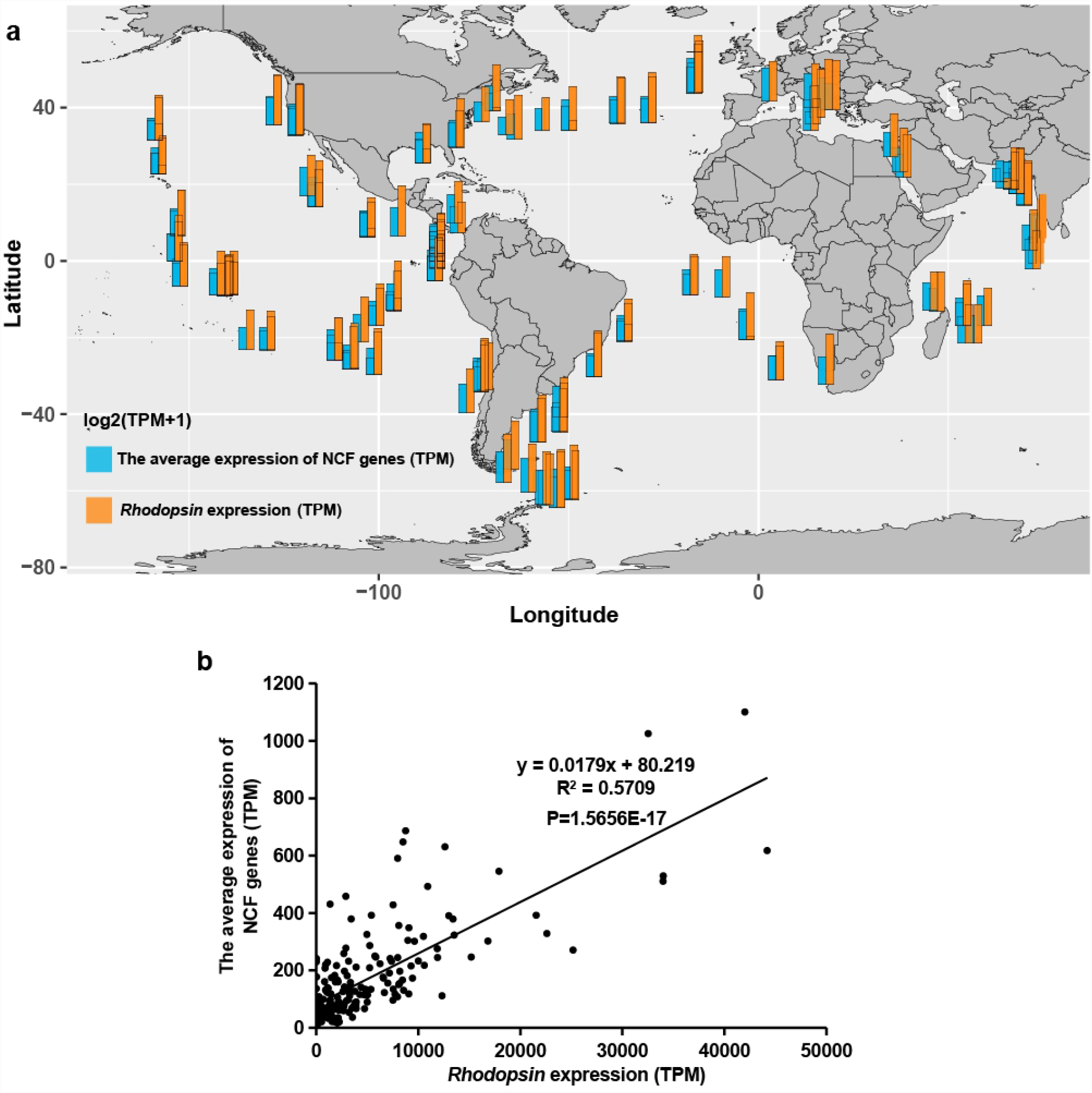
Correlation between the relative activity of non-Calvin carbon fixation (NCF) and the expression of rhodopsin in Flavobacteriales in the global ocean. a, The atlas of NCF gene expression (log2(TPM+1)) and rhodopsin gene expression. b, Linear correlation between the expression level of Flavobacteriales NCF and rhodopsin.

## Discussion

In this study, we used whole-assemblage metatranscriptome sequencing to be able to compare prokaryotes with eukaryotes for common metabolic profiles or physiological functions. Furthermore, using RNA quantity-based calibration, data from separately sequenced size fractions from the same water sample could be integrated to reconstruct assemblage gene expression profile. This allows metabolic estimation of contribution of each size group to the total community in any physiological functions represented by genes expressed in both size groups e.g. free-living bacteria (in the 0.2-3 μm fraction) versus particle-associated bacteria (in the 3-200 μm fraction) to non-Calvin carbon fixation. Using this integrative approach, we obtained novel insights into relative contributions of different lineages to Calvin and non-Calvin carbon fixation, environmental effects, and a potential energy source.

### Calvin carbon fixation by phytoplankton and microbes

For CCF, the relative activity of CCF in the continental shelf area was higher than that in the slope area (Figure 2). This probably because nutrient content is higher at the shelf as it receives greater influences from various nutrient inputs than slope [20, 26]. Indeed, the CCF activity of the major CCF groups was in most cases positively correlated with nutrient contents (Figure S2).

At the supergroup level, CCF potential was mainly contributed by Bacillariophyta, Cyanophyta, Chlorophyta, Haptophyta and O_Stramenopiles in our samples overall. Northern SCS summer phytoplankton communities have been found dominated by Cyanobacteria, Bacillariophyta, Haptophyta and Chlorophyta [27, 28]. This is similar to the taxonomic profile of CCF plankton in our study. The CCF contribution of plankton was, at least in part, correlated with their abundance, as is often found to be the case [29, 30]. In the photic zone, the contribution of Bacillariophyta and Dinophyta to the total CCF of plankton was higher in the shelf area than in the slope area and higher at greater depths. On the contrary, the contribution of Cyanobacteria to the whole plankton’s CCF in the shelf area was lower than that in the slope area and decreased with the increase of depth. This trend is consistent with the general understanding that cyanobacteria are predominant primary producers in the major ocean basins (offshore) whereas diatoms and dinoflagellates dominate nearshore waters, due to contrasting nutrient availabilities.

The major CCF contributors at the level of order were generally similar between the shelf and slope areas, including Bacillariales, Pelagomonadales, Prymnesiales, Mamiellales, and Naciculales, interestingly with no cyanobacterial order, suggesting cyanobacterial contributors to CCF were more taxonomically spread out. Despite the shelf vs slope similarity, the CCF activity of plankton in different orders varied with geographical location and depth. Bacillariales was most active in shelf DCM layer, Mamiellales most active in slope DCM layer, and Dinophysales was most active in the SUR layer. This pattern might reflect niche differentiation among these orders, probably having to do with their differential potential to acquire light energy and nutrients. For instance, diatoms favor high inorganic nutrients [31], whereas Mamiellales species prefer to be distributed in the subsurface layer with high chl *a* concentration [32]. Dinoflagellates are more competitive in nutrient poor environments as they are versatile in scavenging nutrients and they possess proton pump rhodopsin for efficiently harvesting light energy [1], which might help prevents photoinhibition that often occurs on sea surface phytoplankton [33]. Indeed, dinoflagellates were actively expressing rhodopsin (data not shown) and phagotrophy genes in the present study.

### Non-Calvin carbon fixation by microbes

NCF has long been suggested but not become a focused topic of research. A large number of prokaryotes in the ocean have engaged in NCF through at least five pathways, including Reductive Citric Acid cycle (Arnon-Buchanan cycle), reductive Acetyl-Coa Pathway (Wood-Ljungdahl pathway), 3-hydroxyPropionate bicycle, hydroxyPropionate-hydroxybutyrate cycle, and Dicarboxylate-hydroxybutyrate cycle [34]. More recently, NCF was increasingly reported for marine bacterial isolates [12, 13, 35]. However, lineage-specific activities of NCF and their contribution to total carbon fixation are still poorly understood. In the present study, all the five NCF pathways were represented in the whole-assemblage metatranscriptomic dataset, indicating that there were at least five different ways of bacterial NCF in the euphotic zone of the northern South China Sea. Based on the average expression level of complete pathway genes, we found that Flavobacteriales, Alteromonadales, Oceanospirillales and Pelagibacterales were the dominant NCF contributors in the northern SCS for NCF (Figure 5). Generally, prokaryotes in small-sized samples are free-living (FL), while prokaryotes in large-sized samples are particle-associated (PA). Studies in multiple sea areas have shown that the diversity and abundance of PA bacteria are generally higher than that of FL bacteria [36, 37]. In this study, FL and PA prokaryotes contributed nearly equally except in C6_DCM and C9_SUR, where PA prokaryotes contribution was higher than FL prokaryotes. It is rather striking that the PA prokaryotes are potentially equally or more important in non-Calvin carbon fixation.

### Lineage-differential effects of environmental factors on carbon fixation

Marine Calvin carbon fixation is well known to be affected by environmental factors such as light cycle and light intensity [38, 39], salinity [40], temperature [41], N-, P-, and Si-nutrients [42–44], and trace metals [45]. However, the effects differ between species, and it is challenging to investigate lineage-differential effects in a natural community *in situ*. In this study, we examined the relationship between the lineage-specific carbon fixation potential (gene expression) and environmental factors at supergroup and order levels and found differential responses of different lineages to environmental factors. The proportion of CCF contributed by cyanobacteria was negatively correlated with the phosphorus concentration (Figures 2, 3), which was probably because the abundance of cyanobacteria is positively affected by the N:P ratio [46]. The dinoflagellate proportion of CCF was positively correlated with temperature, and significantly negatively correlated with salinity (Figures 2, 3). This may be because these dinoflagellates prefer lower salinity and higher temperature conditions [47].

Regarding non-Calvin carbon fixation, the relative contribution of Cytophagales, Nitrospinales, and Oceanospirillales was significantly positively correlated with salinity. This may indicate that these three orders are more NCF-active in higher salinity environment than other major orders of NCF contributors. In contrast, some orders of bacteria did not show a clear trend in NCF in relationship to environmental factors. This might be because NCF potential of different species from the same order might responded differently to environmental factors. When species coverage of the genome and transcriptome databases increases in the future, the relationship between carbon fixation activity and environmental factors can be studied at a finer taxonomic level, and more trends of NCF with environmental factors might emerge.

### Lineage-differential relationships of endocytosis with Calvin carbon fixation

Endocytosis is usually divided into three types: phagocytosis, pinocytosis, and receptor mediated endocytosis due to the different size of the material and the mechanism [48]. It is increasingly recognized that many phytoplankton are mixotrophic (hence named mixoplankton) [49–51]. Diatoms can take in iron ions in seawater through pinocytosis and even capture silicon to support their growth [52–54]. Many dinoflagellates can perform phagocytosis to cope with the lack of nutrients in the environment [16, 55]. Viruses and commensal microorganisms enter cells of protists through the endocytic pathway of the protistan hosts [56]. Therefore, the endocytosis of these CCF microorganisms may also affect their photosynthesis and carbon fixation. In this study, we analyzed the relationship between photosynthesis and endocytosis of Bacillariophyta, Dinophyta, Haptophyta and Chlorophyta. The results showed that there was a positive correlation between the core genes of endocytosis and the core genes of photosynthesis in Bacillariophyta, Haptophyta and Chlorophyta (Figure 4). This suggests that these diatoms might be taking in iron and other nutrients through pinocytosis to support their photosynthesis and growth [53]. Besides, through phagocytosis the haptophytes and chlorophytes might be obtaining essential growth factors to promote growth of the CCF population [57]. In sharp contrast, a negative correlation was found in dinoflagellates between the expression of phagocytosis core genes and that of photosynthesis core genes (Figure 4). This is probably because many dinoflagellates are mixotrophic and use phagotrophy to compensate low photosynthesis during nutrient limitation.

### Rhodopsin as potential energy source for non-calvin carbon fixation in bacteria

As discussed earlier, studies in recent years have revealed that many prokaryotes in the ocean are engaged in carbon fixation activities through five non-calvin carbon fixation pathways [34]. However, the source of energy to support NCF remains elusive. While there is chemical energy from organic carbon metabolism or the energy from the oxidation of inorganic compounds [58], another potential source is light energy harnessed by proton pump rhodopsin (PR). This structurally very simple and hence highly efficient machinery includes all-trans retinal coupled with an opsin protein and absorbs light energy from the blue-green waveband to pump protons outside of cell membrane, thus creating a proton gradient to drive synthesis of ATP [59, 60]. PR is therefore a photosystem-independent solar energy converting mechanism. PR is widespread in marine bacteria, in which the energy harnessed facilitates survival and growth of the bacteria in nutrient-limited environments [61, 62]. Therefore, we analyzed the rhodopsin expression of the four most important NCF bacterial orders, Flavobacteriales, Alteromonadales, Pelagibacterales and Rhodobacterales. Results showed that the four orders with the highest NCF contribution were also the four orders with the highest rhodopsin gene expression, and the rhodopsin expression level of these four orders was significantly positively correlated with their NCF gene expression (Figure 7). To explore whether the same correlations occur in other parts of the global ocean, we mined the metatranscriptome database of the Tara Oceans expedition. Indeed, the same result was found in *Flavobacterium* (Figure 8). Based on all these results, we hypothesize that rhodopsin is a potential energy source for NCF in some lineages of NCF-active bacteria. Further investigation is warranted to examine this hypothesis.

## Conclusion

There have been several large-scale studies using meta-omics to understand metabolic performance of microorganisms in the ocean [8], but few have dealt with lineage-specific carbon fixation, compared calvin carbon fixation (CCF) with non-calvin carbon fixation (NCF), and their relationships with environmental effects. Even less effort has been focused on the South China Sea, a large continental marginal sea. This first attempt to catalog functional genes in China Seas documented over 4.4 million unigenes. This will be a useful baseline data for future research on plankton metabolic pathways *in situ* in this area. Previous metatranscriptomic studies have tended to focus on either eukaryote or prokaryote, and our whole-assemblage metatranscriptomics approach include both groups. This allows us to make a comparative analysis of contributions of prokaryotes and eukaryotes to common physiological processes at the level of gene expression, and better understand the interaction between eukaryotes and prokaryotes. At the transcriptome level, we have assessed the contribution of the major carbon-fixing plankters in different water layers in the shelf and slope areas of the northern South China Sea. We found that Bacillariales, Pelagomonadales, Prymnesiles, Mamiellales and Naviculales were the major contributors of CCF, while Flavobacteriales, Alteromonadales, Oceanospirillales and Rhodobacterales were most active in NCF. CCF and NCF capacities were higher at the shelf and differentially affected by environmental factors depending on lineages. Besides, endocytosis can promote CCF in diatoms, chlorophytes, and haptophytes, while it may complement CCF in dinoflagellates. NCF in Flavobacteriales, Alteromonadales, Pelagibacterales and Rhodobacterales appears to be energetically supported by proteorhodopsin. Moreover, the bioinformatics data obtained in this study and the research methods adopted lay a foundation for further studies on lineage-specific plankton metabolism.

## Materials and Methods

### Field sampling

During a research cruise in the period of 6-12 August 2016, 20 samples were collected at C6 (117.46°E, 22.13°N) and C9 (117.99°E, 22.69°N) stations from the northern South China Sea (SCS), separated into the picoplankton/nanoplankton (0.2-3μm) and microplankton (3-200μm) organismal size fractions. Stations C6 and C9 are located on the shelf and slope of the northern South China Sea, respectively. Sampling targeted three depth layers, which encompassed distinct physicochemical conditions: surface (SUR), deep chlorophyll maximum (DCM) and bottom of photic zone (BOT) layers. For each sample, 40-50L seawater from more than three CTD rosettes (one CTD rosette contain 12L seawater) was first filtered through 200μm blotting cloth to remove large plankton. Then the seawater was serially filtered through two size fractions (0.2-3 μm and 3-200 μm) using 142mm polycarbonate membranes (Millipore, Billerica, MA, USA), each sample taking about 20min. Finally, the sample-carrying filters were divided into four quarters: one being stored in a 2ml tube containing lysis buffer (0.1M EDTA and 1% SDS) for DNA work and were in Trizol reagent (Molecular Research Center, Inc, USA) for RNA work. All samples were immediately frozen in liquid nitrogen onboard until processing after return to the laboratory. Two replicate samples were collected from each condition, a total of 20 samples were collected (Figure 1).

### Measurements of physical and chemical parameters

Temperature, salinity, and depth were measured using a CTD profiler (SBE 17plus V2, Sea-Bird Scientific, United States). A series of nutrients (NO_3_ ^-^+NO_2_ ^-^, PO_4_ ^3-^, SiO_3_^2-^) were measured using continuous flow analyzer, all measurements were performed on triplicate samples. The light intensity of each water layer was calculated based on remote sensing data and a reported computational algorithm [63].

### Sample processing and metatranscriptomic sequencing

RNA was extracted as previously described [6]. Shortly, each of the samples was mixed with a 1:1 mixture of 0.5 mm and 0.1 mm-diameter zirconia/silica beads (Biospec, USA), and beat at the rate of 6 m/s on a FastPrep-24 bead mill (MP Biomedicals, USA) for three times to ensure complete cell breakage. RNA was extracted following the TRI Reagent protocol coupled with the Direct-zol™ RNA columns, essentially as reported previously [1]. DNase 1 was used to remove DNA from the total RNA. RNA concentration was measured using a NanoDrop ND-2000 Spectrophotometer, and integrity was assessed using RNA 6000 Nano LabChip Kit in microcapillary electrophoresis (Agilent 2100 Bioanalyzer, Agilent Technologies, Australia). Samples with the RNA integrity number (RIN) ≥ 6.0 were used for metatranscriptome sequencing. One μg RNA from each sample was subject to ribosomal RNA removal using a Ribo-Zero rRNA Removal Kit (Plant Leaf) and a Ribo-Zero rRNA Removal Kit (Plant Seed/Root) (Illumina, San Diego, CA, United States) for whole-assemblage (prokaryotes + eukaryotes) metatranscriptome (WAM) sequencing. mRNA (i. e. rRNA depleted RNA) was then fragmented with First Strand Synthesis Reaction Buffer and Random Primer Mix (2×) at 94°C for 10 min, and first strand cDNA was synthesized using ProtoScript II Reverse Transcriptase and the second-strand cDNA was synthesized using Second Strand Synthesis Enzyme Mix. The double-stranded cDNA was purified, end repaired, and ligated to adaptors. Fragments of about 400 bp (with the approximate insert size of 250 bp) were selected and sequenced on Illumina HiSeq 4000 instrument (Illumina, San Diego, CA, United States). Each sample had two or three technical replicates, and totally produced 881Gb raw data.

### Transcriptome analysis and gene expression quantification

Raw reads were processed by removing adaptors, reads with >5% ambiguous bases (N), low quality reads (>20% bases with quality value < 20). For the remaining clean reads, we used Trinity to perform *de novo* assembly, then used Tgicl to cluster transcripts to unigenes. The unigene sets from all samples were clustered again to generate the final unigene set (Unigene) for downstream analysis. We used BLASTN [64] for NT annotation, blastx [64] or Di for NR, KOG, KEGG and Swissprot annotation, blast2go [65] and NR annotation results for GO annotation, and interprocan5 [66] for interpro annotation. Bowtie2 [67] was used to match clean reads to Unigene, and then Salmon v0.9.1 [68] was used to calculate gene expression levels of each sample. In the subsequent analysis, we excluded unigenes whose TPM (Transcripts Per Kilobase of exon model per Million mapped reads) was less than 0.1 across all 20 samples.

### Identification of co-highly expressed genes and expression of key genes in different samples

To obtain a robust actively expressed gene set, highly expressed genes (HEGs) were identified in each of the samples. HEGs were defined as genes with TPM in top 25% in each sample. Common HEGs in C6 and C9 stations were used to conduct KEGG enrichment to obtain active functional pathways. Key genes in carbon fixation were identified from the WAM dataset based on functional annotations. The expression contributions of these genes were visualized using Circos (http://mkweb.bcgsc.ca/tableviewer/visualize/).

### Correlation of expressions of genes in different physiological functions

We analyzed correlations for genes with non-zero expression in at least six out of our 20 samples. Correlation between the expression of carbon fixation core genes and endocytosis core genes were visualized in R using packages ggplot [69]. Pearson correlation was analyzed between the expression of non-Calvin carbon fixation and that of proteorhodopsin in bacterial lineages in which proteorhodopsin expression was high.

### Quantification of the relative activity of carbon fixation

Based on the annotations of the KO database, we used the TPM value of *RuBisCO* gene to represent the CCF activity of the lineage (supergroup). Besides, we counted the sum of the TPM values of all unigenes on the NCF pathway for individual lineages of interest and calculated the average TPM value of each gene in the pathway. The average value was used to represent the NCF activity of the lineage (supergroup). The greater the TPM value, the greater the activity of carbon fixation.

### Calibration for size fraction dataset to enable comparison

Large (3-200 µm) and small (0.2-3 µm) size fractions were sequenced separately, making it impossible to directly compare contributions of these two size groups to the community in various metabolic processes at the transcriptional level. To overcome the problem, we multiplied TPM of a functional gene in a size fraction with the total RNA extracted from the size fraction sample, then divided the product by the total RNA extracted from both size fractions from the same volume of water sample, i.e. TPM_small_ * RNA_small_ per L/(RNA_small_ per L + RNA_large_ per L) and TPM_large_ * RNA_large_ per L/(RNA_small_ per L + RNA_large_ per L). These allow estimation of contribution of a small-sized plankton or a large-sized plankton to the whole-assemblage in the function represented by the gene (e.g. *RuBisCO*).

### Nucleotide Sequence Accession Numbers

The data that support the findings of this study have been deposited in NCBI database (http://www.ncbi.nlm.nih.gov/) under the BioProject number PRJNA729123 and CNGB Sequence Archive (CNSA) of China National GeneBank DataBase (CNGBdb) (https://db.cngb.org/cnsa/) under the accession number CNP0001483.

## Author contribution

Senjie Lin designed the research. Tangcheng Li and Cong Wang collected samples. Hongfei Li performed the experiments. Hongfei Li, Jianwei Chen, Liying Yu, Guangyi Fan, Ling Li, Tangcheng Li, Huatao Yuan, Jingtian Wang, Cong Wang and Senjie Lin analyzed the transcriptomic data. Hongfei Li and Senjie Lin wrote the manuscript. All authors read and approved the manuscript.

## Acknowledgment

We wish to thank Bangqin Huang, Xin Liu, Yanping Zhong, and Chentao Guo of Xiamen university for help with sampling and logistic support. We are also indebted to the crew and participants of the Yanping 2 research cruise for assistance in sampling. In addition, we thank Xin Lin for participating in various discussions of the study. This work was supported by National Key Research and Development Program of China grant 2016YFA0601202 and the Marine S&T Fund of Shandong Province for Pilot National Laboratory for Marine Science and Technology (Qingdao) (No.2018SDKJ0406-3).

