## supplementary information for "Differential CO_2_-fixation potentials and supporting roles of phagotrophy and proton pump among plankton lineages in a subtropical marginal sea"

**Table S1. The physical and chemical parameters**

| | C6S_1 | C6S_2 | C6D_1 | C6D_2 | C9S_1 | C9S_2 | C9D_1 | C9D_2 | C9B_1 | C9B_2 |
|---|---|---|---|---|---|---|---|---|---|---|
| $NO_2^-$+$NO_3^-$ (μm) | 0.3985 | 0.272 | 1.2915 | 0.281 | 0.607 | 0.156 | 0.584 | 2.291 | 9.108 | 8.246 |
| $PO_4^-$ (μm) | 0.058 | 0.053 | 0.538 | 0.087 | 0.011 | 0.021 | 0.131 | 0.242 | 0.717 | 0.648 |
| $SiO_3^{2-}$ (μm) | 3.0145 | 2.065 | 2.7505 | 1.557 | 1.527 | 1.497 | 2.702 | 3.61 | 8.641 | 7.769 |
| Temperature (°C) | 29.26 | 28.5935 | 27.0803 | 27.0055 | 30.2508 | 29.8893 | 24.5048 | 26.2942 | 18.61 | 18.9146 |
| Salinity | 33.62 | 34.2943 | 34.1409 | 34.4909 | 34.0557 | 33.7776 | 34.6454 | 34.7504 | 34.7678 | 34.7839 |
| Depth (m) | 3 | 3 | 35 | 35 | 3 | 3 | 75 | 75 | 150 | 150 |
| PAR (μmol $m^{-2}$ $s^{-1}$) | 55.8697 | 55.8697 | 4.007 | 4.007 | 59.0434 | 59.0434 | 0.7454 | 0.7454 | 0.0221 | 0.0221 |

**Table S2. Statistics of unigene annotation**

| Annotation | Annotated No. | Annotated% |
|---|---|---|
| **Nr** | 1,532,571 | 34.44% |
| **Nt** | 431,723 | 9.70% |
| **Swissprot** | 808,578 | 18.17% |
| **KEGG** | 666,817 | 14.99% |
| **GO** | 689,985 | 15.51% |
| **Overall** | 1,656,453 | 37.23% |

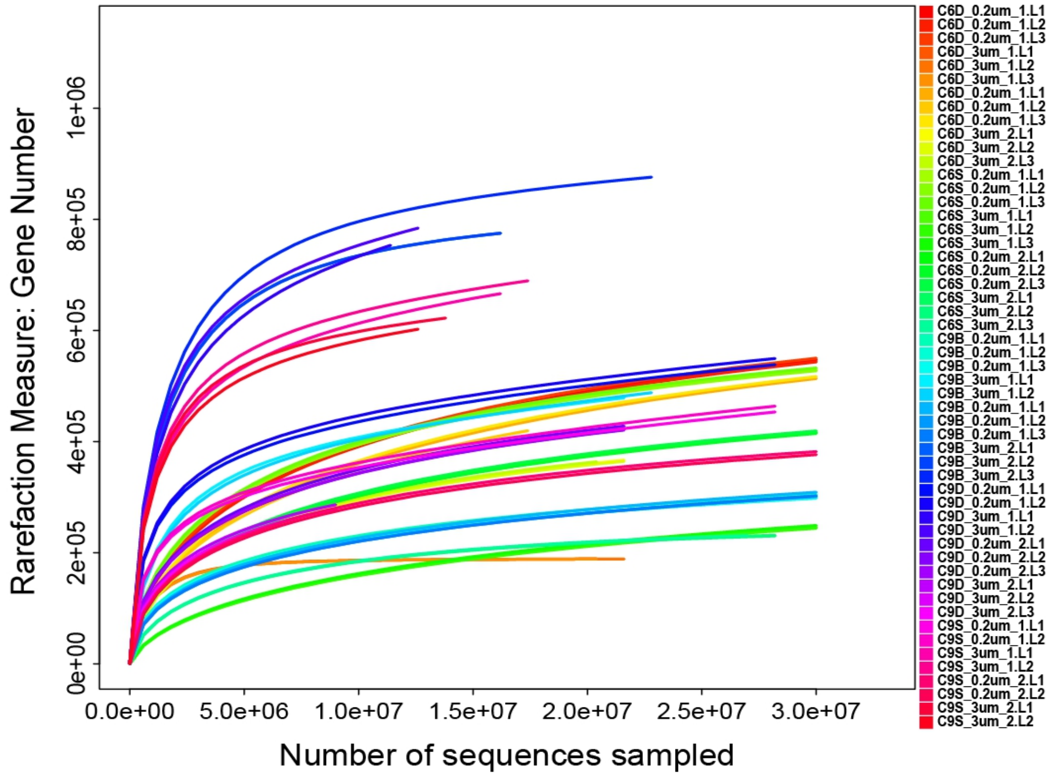


**Figure S1. Rarefaction curves of detected genes from metatranscriptome sequencing output showing that sequencing scale was nearly saturated.** Each plankton sample has 2 to 3 sequencing replicates.

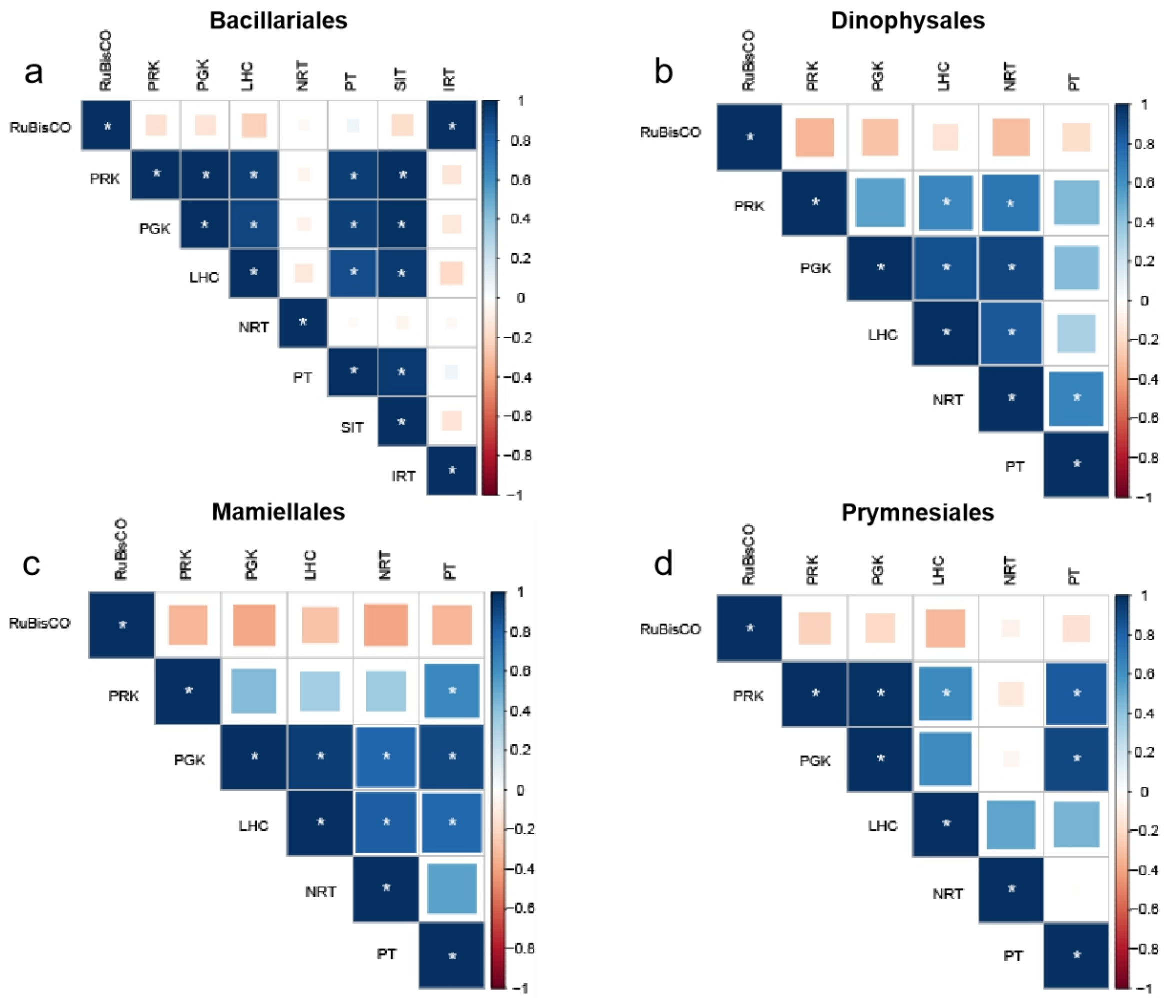


**Figure S2. The relationship between Calvin carbon fixation (CCF) activity.** PRK: phosphoribulokinase, PGK: phosphoglycerate kinase, LHC: light-harvesting complex-like protein, NRT: nitrate/nitrite transporter, PT: phosphate transporter, SIT: Silicon transporter, IRT: Iron transporter. * means $P < 0.05$

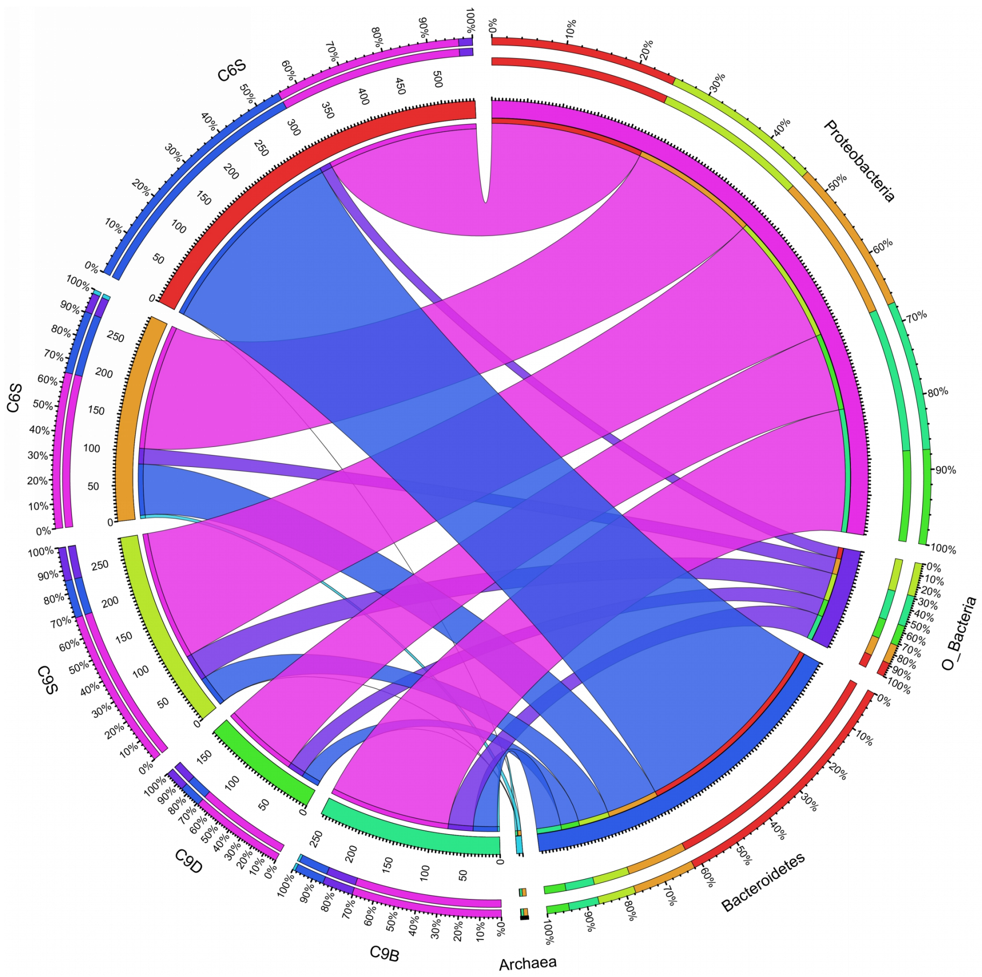


**Figure S3. The relative activity of non-Calvin carbon fixation (NCF) in different water layers and the proportion of contribution by major NCF lineages.** The inner circle of numbers are the average expression of NCF pathway genes in the specified station and depth, as proxy of relative activity of NCF.

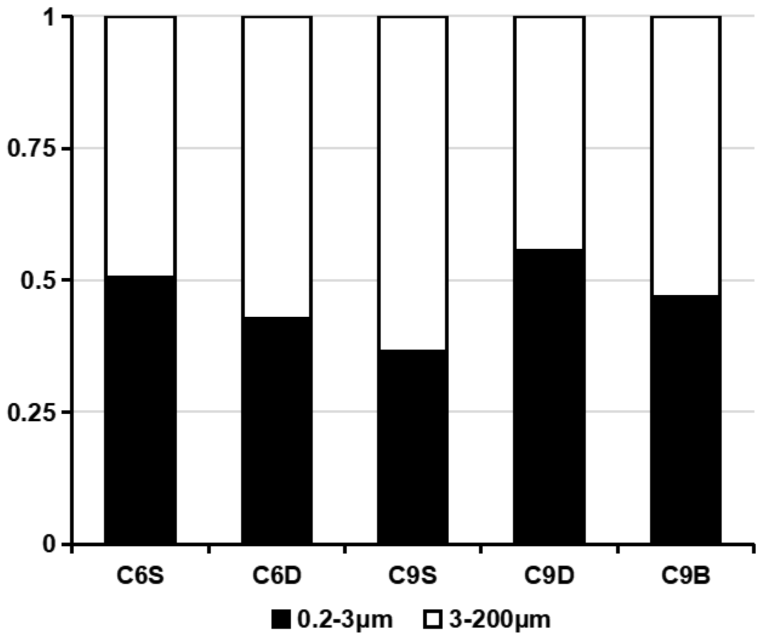


**Figure S4. Relative contribution (%) of non-Calvin carbon fixation (NCF) by two size fractions.**
